# Changes in cuticle composition co-regulate drought and herbicide resistance in horseweed (*Erigeron canadensis*)

**DOI:** 10.64898/2026.06.16.732734

**Authors:** Michael Ozolins, Ahmet Tansel Serim, Mohit Mahey, Sara Álvarez-Rodríguez, Eric L. Patterson

**Affiliations:** Department of Plant, Soil and Microbial Sciences, Michigan State University, East Lansing, MI, USA; Department of Plant Protection, Bilecik Seyh Edebali University, Bilecik, Turkey

**Author notes:** Author for correspondence: Eric L. Patterson. Michael Ozolins, Ahmet Tansel Serim, Mohit Mahey, Sara Álvarez-Rodríguez.

**Keywords:** Cuticle, Drought stress, Glyphosate, Herbicide absorption, Herbicide resistance, Triterpenoids

## Abstract

Horseweed (*Erigeron canadensis*) is a widely distributed annual weed that can cause significant yield losses if not properly controlled. Its phenotypic plasticity allows it to rapidly acclimate to new environmental conditions, such as drought and herbicides, such as glyphosate, with the potential for cross stress acclimatization. The objectives of this research were to uncover the physiological and genetic effects at the intersection of drought stress and glyphosate resistance. To this end, we performed greenhouse dose response experiments, RNAseq, ^14C^ glyphosate absorption and translocation, and cuticular lipid profiling via GC/MS. Greenhouse dose-response experiments revealed that, after drought stress, there was a 2.5-3.7 fold reduction in glyphosate sensitivity via a significant reduction in glyphosate absorption, regardless if the starting population was resistant or susceptible to the field use rate already. Cuticular waxes were collected from each population with and without drought stress and were analyzed via GC/MS. When comparing total wax loads of plants grown under WW and DS conditions, we found that drought stress significantly increased total wax loads for all three populations. Additionally drought stress substantial increases the proportion of triterpenoids in the cuticle. By RNAseq, we found serval triterpenoid biosynthesis genes upregulated after drought, which likely drive the changes in cuticle composition and ultimately increased glyphosate resistance following drought. Ultimately, understanding how drought impacts glyphosate resistance is critical for maintaining optimal weed control in the changing climate.

**Highlight:** Drought stress induces changes to cuticle composition and gene expression that reduce glyphosate absorption, thereby increasing horseweed’s ability to survive glyphosate application.

## Introduction

Herbicide-resistant weed populations evolve from selection pressures imposed by intensive, repetitive herbicide applications (Norsworthy *et al*., 2012). As we increasingly rely upon herbicides, weeds that evolve even moderate resistance gain a significant competitive advantage and outcompete susceptible counterparts as well as crops. Horseweed, a ubiquitous broadleaf weed found across North America, has emerged as a particularly problematic species, with numerous biotypes demonstrating resistance to a wide range of herbicide mechanisms of action, including glyphosate. Glyphosate was a once-dominant herbicide that is losing its supremacy due to widespread resistance evolution in almost every major weed species. This has led to major concerns about our ability to sustainably control weeds in many row crop systems (Owen, 2016). Horseweed was the first broadleaf weed species reported to evolve glyphosate resistance (GR) (VanGessel, 2001) and may be ranked at the top of the most troublesome weed list in large part due to herbicide resistance to chemistries from HRAC groups 2, 5, 9, and 22 (Hanson *et al*., 2009) (Heap, 2024).

The ability for horseweed to evolve and spread resistance is due to several unique traits. Individual horseweed plants produce hundreds if not thousands of lightweight, wind-dispersed seeds (Liu *et al*., 2022) that continuously germinate throughout the growing season, forming both summer and winter annual populations (Davis *et al*., 2009). Strategies to mitigate the evolution of glyphosate resistance in horseweed require a deep understanding of the various mechanisms it employs for resistance. Shikimic acid assimilation tests have found that many of the glyphosate-resistant biotypes exhibit altered glyphosate metabolism, suggesting metabolic resistance mechanisms rather than a target-site mutation in the target protein 5-enolpyruvylshikimate-3-phosphate synthase (EPSPS) (Hanson *et al*., 2009). Additionally, Pro-106-Ser mutations in *EPSPS* have been found in some populations that confer high levels of resistance to glyphosate (Beres *et al*., 2020; Sulzback *et al*., 2025). Furthermore, differences in glyphosate translocation are also often cited as resistance mechanisms with the M10, M11, M7, and P3 *ABC transporter* genes being implicated (Tani *et al*., 2016). These biotypes may be sequestering glyphosate into the vacuole, effectively isolating it from the target site and preventing it from exerting its herbicidal effects, thereby leading to increased resistance (Ge *et al*., 2010).

Regardless of the mechanism, herbicide resistance can be further exacerbated by adaptations and physiological changes caused by other environmental factors, including drought, which can significantly influence the growth, development, and stress responses of weeds. It is often unclear how such stressors shape the evolution and proliferation of herbicide resistance in problematic weeds like horseweed (Norsworthy *et al*., 2012; Kraehmer *et al*., 2014; Owen, 2016). Many resistance mechanisms also seem to enhance the ability of these plants to withstand adverse environmental conditions such as drought, salinity, high/low temperature, and heavy metal toxicity; either through reshaping plant stress response mechanisms, defense mechanisms, or physiological barriers between the plant and the environment (Pandian *et al*., 2020). Co-adaptation of weeds to drought and herbicides complicate the already complex task of managing weeds, particularly in the face of a changing climate that may increase the frequency and intensity of drought events (Weller *et al*., 2019).

Previous research supports a close relationship between herbicide resistance and the drought stress responses of weeds (Mohammad *et al*., 2022). Mohammad et al. (2022) showed that drought-stressed *Alopecurus myosuroides* leads to increased herbicide resistance, and that this phenotype can be inherited possibly via a transient, epigenetic-based, mechanism. Another study conducted on *Eragrostis plana* indicated that this species had transgenerational memory triggered by drought stress and low rates of glyphosate (Fipke *et al*., 2022). Benedetti et al., 2020 reported similar results for one of the most troublesome weeds, *Echinochloa colona*, under drought stress.

In all these cases, the shift in drought and resistance phenotypes was observed, but the specific mechanisms are still unexplored or vague. Although horseweed is an important arable weed in row crops, the potential impact of drought stress on the development and spread of herbicide resistance in horseweed has not been extensively explored. Kanatas et al., 2023 indicated that GR horseweed biotypes had higher germination rates than glyphosate susceptible (GS) horseweed biotypes under drought stress. (Mojzes *et al*., 2020) showed that this weed can grow more vigorously in moderate drought plots than in control and watered plots. Therefore, understanding the mechanism by which drought tolerance and herbicide resistance in horseweed is crucial for devising effective and sustainable weed management strategies.

This study aims to understand the interactions of drought stress and herbicide resistance in horseweed, taking a mechanistic approach to understand how these two seemingly unrelated intersect and may influence each other. Specifically, this research explores how glyphosate resistant and susceptible biotypes respond to drought and glyphosate separately and in succession, as well as the actual physiological and molecular mechanisms that enable horseweed to maintain or enhance herbicide resistance after drought stress.

## Materials and methods

### Seedling Preparation

Seedlings from one susceptible biotype (Biotype 125) and three resistant biotypes (Biotypes 119, 127, and 135) were used to investigate the impact of drought stress on the physiology of horseweed. These populations were originally collected from a previous survey study (Sulzback *et al*., 2025). Rectangular pots (30x30 cm) were filled with potting soil (Suremix), and seeds were evenly spread on the surface. The pots were watered regularly as needed with lightly fertilized water (NPK, 15-07-25, ICL Specialty Fertilizers, Summerville, South Carolina, USA). When the horseweed seedlings reached the 2-3 true leaf stage, they were transplanted into new pots (12x12 cm) filled with fresh potting same soil. The seedlings were grown in the Michigan State University greenhouse, where conditions were set to a 14-hour photoperiod with day/night temperatures of 26-21°C±1.0 and 23-18°C±1.0, respectively, for 3-4 weeks until they reached the rosette stage during the autumn and winter seasons. The seedlings were irrigated and fertilized every other day.

### Baseline Glyphosate Resistance Evaluation Study

Dose-response bioassays were conducted in a greenhouse to determine baseline glyphosate sensitivity in all four horseweed biotypes. When the seedlings reached the rosette stage, they were sprayed with glyphosate ammonium salt + ammonium sulfate (0.1% v/v) in a spray cabinet calibrated to deliver 20 gallons per acre (1 mph and 30 psi). Glyphosate was applied at rates of 0.63, 1.26, 2.52, 5.04, 10.08, and 20.16 kg ae ha[¹, equivalent to the recommended use rate, where 1.26 kg ae ha[^1^ corresponds to a typical field rate (1x). After application, the seedlings were placed in a greenhouse for 21 days. Following this period, the seedlings were cut at ground level to measure dry weight and stored in a drying oven at 60°C for 72 hours. The experimental design was a randomized complete block with four replications. The entire dose response was repeated once. Dose-response data were evaluated using a nonlinear regression model in R statistical software (RStudio Team, 2023). The DRC package in R was used to determine the dose-response curve and parameters using a three-parameter log-logistic model (Equation 1).

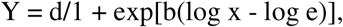

Where:

Y represents seedling dry matter at herbicide treatment rate x b, d, and GR50 represent the slope, upper limit, and herbicide rate that reduces seedling dry matter by 50%, respectively

Horseweed seedlings from Biotypes 125, 119, 127, and 135 were classified as susceptible **(S)** (Population 125), resistant **(R)** (Populations 119, and 127), and highly resistant **(RR)** (Population 135) to glyphosate, respectively, based on the dose-response assay. Biotypes 119 and 127 exhibited similar GR_50_ values, so Biotype 119 was selected to represent the resistant biotype for the remainder of this research.

### Drought x Glyphosate Study

Fully grown seedlings from the three biotypes (S - population 125, R – population 119, and RR - population 135) were divided into two groups (Fig 1). Group 1 was subjected to drought stress. Drought stress was conducted following the protocol described by Dos Santos et al. (2015), recognizing that horseweed is highly tolerant to drought stress. That is to say, the plants were subjected to drought conditions by adjusting to half of the total water-holding capacity (CW) of the pot for 14 days. The total water-holding capacity of the pot was calculated using the following equation:

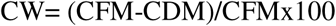

**Figure 1.**
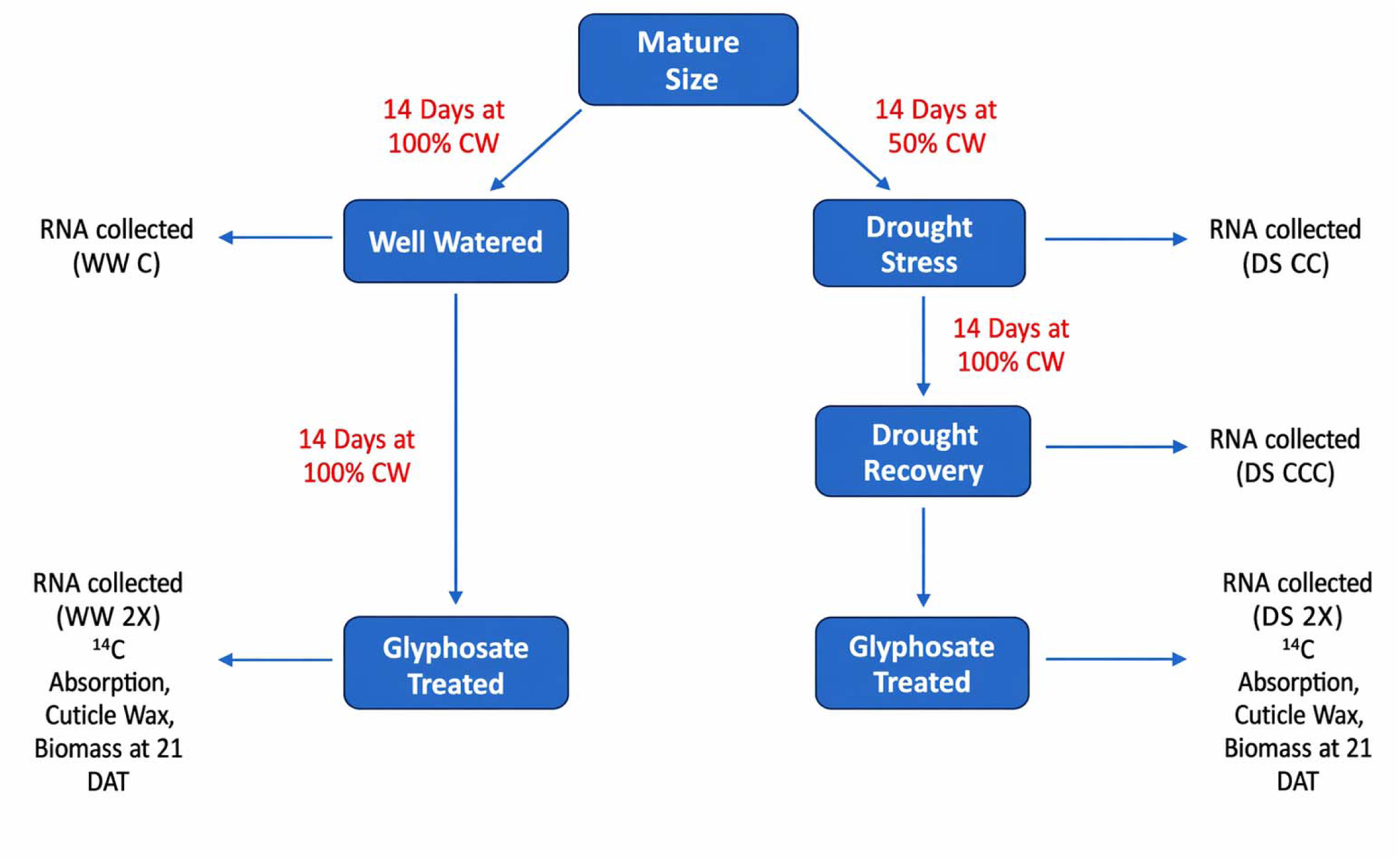
Experimental design and treatment structure. Plants were grown to a mature stage and then assigned to either a well-watered control or a drought stress treatment. A subset of drought-stressed plants was subsequently rewatered to allow drought recovery. Glyphosate was applied to plants under well-watered conditions and to plants following drought recovery, enabling comparisons among well-watered, drought-stressed, and drought-recovered plants with respect to physiological responses and glyphosate uptake and effects. WW C refers to well watered controls, WW 2X refers to well plus glyphosate, DS CC refers to drought stress, DS CCC refers to drought plus recovery, DS 2X refers to drought stress plus glyphosate. CW: Total water-holding capacity of the pot

Where:

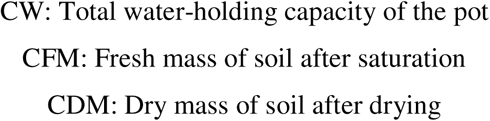

These plants were then allowed to recover for 14 days in well-watered conditions. Group 2 remained well-watered, as described earlier. Dose-response bioassays were repeated to determine the efficacy of drought stress on glyphosate resistance and sensitivity horseweed biotypes using the method aforementioned. To understand the intersection between drought stress and glyphosate resistance in horseweed populations in terms of transcriptomic and metabolomics approaches, an assay was conducted using well-watered and drought-stressed horseweed seedlings. Glyphosate was then applied to both Group 1 after drought and recovery as well as Group 2 at 0X (i.e. untreated) and 2X rates. The experiment followed a completely randomized factorial design (2 × 3 × 2) with four replications. The factors included water regime (well-watered vs. drought-stressed), biotype (susceptible, resistant, and highly resistant), and glyphosate rate (0× and 2×). Treatments were 1) well-watered, 2) glyphosate-treated, 3) drought-stressed, 4) drought-stressed and recovered, and 5) drought stressed-recovered-glyphosate treated plants from susceptible, resistant and highly resistant populations.

Leaf tissues were collected from the youngest fully developed leaves of each treated and untreated seedling 1 DAT, put in 1.5 ml Eppendorf tubes, and immediately flash frozen in liquid nitrogen. The samples were stored in a freezer at -80°C until extraction.

### RNA extraction

The tissues were ground using plastic sterile pastels, cooled in liquid nitrogen. Total RNA was obtained from the leaf tissue using the Plant RNeasy Mini kit (Qiagen, Valencia, CA, USA) following the manufacturer’s recommendations with small modifications. To help remove polysaccharides and genomic DNA, the PVP was added and DNAse treatment was performed on the RNA extractions. The content and quality of RNA were checked using a NanoDrop 1000 spectrophotometer and Tapestation 4150 (Thermo Scientific, Waltham, MA, USA; Agilent Technologies, Inc.) and sent to Novogene for cDNA library preparation and RNA sequencing.

### Cuticular Wax Analysis

Cuticle surface waxes were extracted from parallel leaf tissue. First 5 μg of the internal standard n-tetracosane (24:0 alkane) was added to each sample and then they were extracted by immersing them in 10 mL of chloroform for 2 minutes. Chloroform was evaporated under a gentle stream of nitrogen gas. Wax extracts were then converted into their trimethylsilyl derivatives by dissolving the samples in 100 μL of N,O-bis-(trimethylsilyl)-trifluoroacetamide (BSTFA) along with 100 μL of pyridine at 65 °C for 16 h. For cuticle chemical identification, the samples were analyzed on an Agilent 5975 mass spectrometer. A 1-10 spilt injection was used with a VF-5ms column (30 m length, 0.25 mm inner diameter, 0.25 μm film thickness). Temperature settings were as follows: inlet 300 °C, oven temperature program was set to 80 °C for 1 min and increased to 325 °C at a rate of 40 °C per minute. The oven temperature was then held at 325 °C for 10 min. The helium flow rate was set at 1 mL per minute. Data were acquired in full-scan mode over a mass range of m/z 50–650 with a solvent delay of 7.0 min. Compounds were identified based on comparison of their mass spectra to authentic standards and NIST library database. Each leaf sample was scanned, and its surface area was measured using ImageJ.

### Absorption and Translocation

The most developed leaf from each group of plants, both well-watered and drought-stressed, was chosen for the application of radiolabeled [^14^C] glyphosate. Each plant was treated with 200,000 DPM of [^14^C] glyphosate. The spotting solution contained [^14^C] glyphosate, unlabeled glyphosate, AMS, and water to achieve the same concentration as a glyphosate application of 1.27 kg ha–1. Each leaf was treated by applying 10 1-uL droplets on the upper leaf surface and then placed in a growth chamber maintained at 25/18°C day/night temperature with a 16-hour photoperiod.

Plants were harvested at 1, 3, 6, 12, 24, and 72 hours after treatment. At harvest, each plant was separated into the treated leaf and the rest of the aboveground tissue. Unabsorbed [^14^C] glyphosate was removed by placing the treated leaf in a 20 ml scintillation vial with 5 ml of a methanol: water (20:80) solution and shaking it for 1 minute. Then, the treated leaf was transferred to a new scintillation vial, and the process was repeated. Each plant part was dried for 48 hours before combustion in a biological sample oxidizer counter (Sample Oxidizer Model 307; PerkinElmer, Boston, MA) for 1 minute. The ^14^CO_2_ released from the biological oxidizer was trapped in 20 ml of scintillation fluid (Carbo-Sorb® E:Permafluor® E+, 1:1 v/v; PerkinElmer, Groningen, Netherlands), and the radioactivity was measured using a liquid scintillation counter (Tricarb 4910TR Liquid Scintillation Analyzer; PerkinElmer, Boston, MA). The radioactivity in the 5-ml leaf wash solution was quantified with the addition of 15 ml of Ultima Gold™ scintillation fluid (PerkinElmer, Groningen, Netherlands).

Glyphosate absorption was calculated as the sum of the total ^14^C in the plant parts divided by the total ^14^C recovered, including the treated leaf wash. The amount of ^14^C present in the leaf wash and the plant sections was considered as total 14C recovered, which averaged >90% of applied [^14^C] glyphosate. Translocation out of the treated leaf was calculated by taking the amount of ^14^C absorbed in the untreated plant parts divided by the total ^14^C absorbed in the plant.

### Sequencing and RNAseq

The extracted RNA was sequenced using paired-end 150bp Illumina hiseq sequencing platform. The reads were cleaned, and trimmed using fastP (Chen *et al*., 2016) using the default filters. The cleaned reads were aligned using HiSAT2 v2.1.0 (Kim *et al*., 2019) to the reference genome of horseweed (Laforest *et al*., 2020). FastQC v0.11.7 (Andrews 2010) along with multiQC (Ewels *et al*., 2016) was used to obtain trimming, filtering, and alignment metrics. Samtools v1.16.1 (Li *et al*., 2009) was used to convert alignment files and for indexing. FeatureCounts v2.0.6 (Liao *et al*., 2014) was used for extracting aligned reads counts. All the commands utilized can be found on GitHub (https://github.com/mohitmahey/RNA_seq_analysis).

### Weighted gene co-expression analysis (WGCNA) –

Gene co-expression analysis was conducted using the WGCNA v1.72-5 R package (Langfelder and Horvath, 2008). The raw counts were normalized and averaged across replicates. The read counts were normalized using the DESeq2 v1.44.0 package (Love *et al*., 2014). A PCA was made with normalized read counts across population and replicates. Hierarchical sample clustering was done using “average” method to find sample outliers. Soft-threshold power estimated from 1 to 24 for network construction by calculating scale-free topology fit index. This was performed to prevent batch effects and create a scale-free network with cutoff threshold of 0.8. The signed adjacency matrix was generated using biweight midcorrelation, and the signed topological overlap matrix was obtained from dissimilarity measures. Genes were organized into clusters through hierarchical clustering, with a minimum module size set at 30 and a merge cut height of 0.25. The dynamic tree-cutting algorithm was then utilized to assign genes into separate modules. GO term enrichment analysis was conducted using topGO v2.56.0. All the analysis was done using R v4.4.0

## Results and Discussion

### Dose response well-watered and drought stress

Under well-watered conditions, the dose of glyphosate required to cause 50% reduction in plant dry weight (GR_50_) was 0.226 kg a.e. ha^-1^ for the susceptible population, 5.03 kg a.e. ha^-1^ for the R population, and 22.31971 kg a.e. ha^-1^ for the RR population (Fig 2a). Under drought conditions there was a significant increase in GR_50_ for all three populations, 0.51 kg a.e. ha^-1^ for the S population, 19.74 kg a.e. ha^-1^ for the R population, and 47.40 kg a.e. ha^-1^ for the RR population (Fig 2a). In summary, after drought stress, there was a 2.5-fold increase in glyphosate resistance for S, 3.7-fold increase in glyphosate resistance for R, and 2.45-fold increase in glyphosate resistance for RR (Table 1). These results indicate that previously experienced drought stress decreases glyphosate sensitivity in individual horseweed plants in both sensitive and resistance populations; with the largest effects appearing in populations already resistant to glyphosate (Fig 2b). This adds to a growing body of evidence that drought stress can increase herbicide resistance both within a single generation (i.e. stress priming) and transgenerationally (Fipke *et al*., 2022). The exact mechanisms that cause this phenomenon are still unclear; however, it has been shown that drought can prime plants for future stress by altering the plants’ metabolism and base biochemistry. For example, drought can highly impact cuticle thickness and composition which would theoretically lead to decreased or altered foliar applied herbicide absorption (Oosterhuis *et al*., 1991) *Glyphosate Absorption and Translocation under Well watered and Stress Primed Conditions*

**Figure 2.**
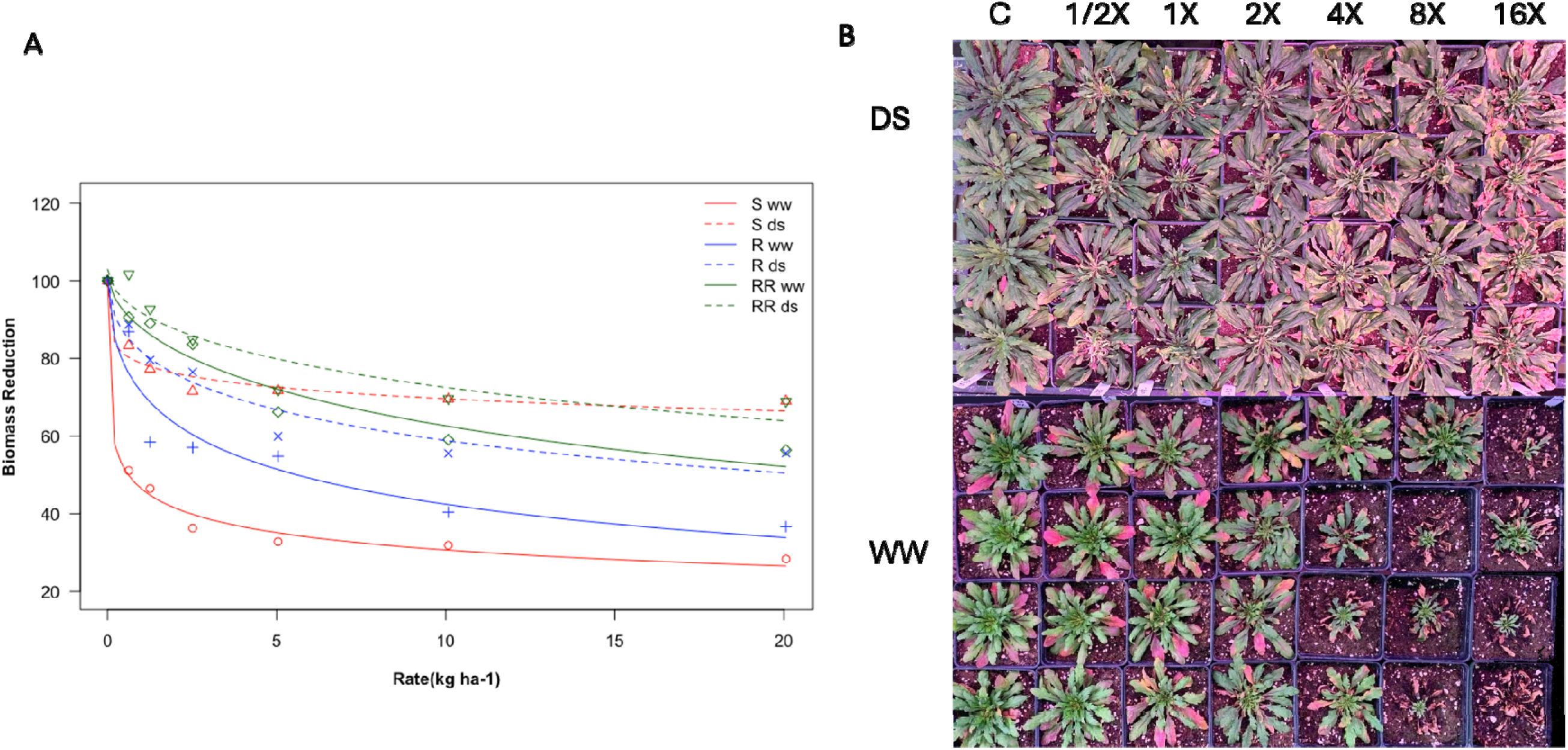
(A) Dose-response assay of *Erigeron canadensis* biotypes RR (highly resistant), R(resistant), and S(susceptible) treated with glyphosate before and after drought stress. Biomass is expressed as a percentage of the nontreated control of each biotype and drought treatment. (B) Photos of dose-response assay with glyphosate before and after drought stress. DS refers to drought stress, WW refers to well water, X refers to field rate (1.26 kg a.i. ha^-1^)

**Table 1.** Dose response analysis of three horseweed (Erigeron canadensis) populations which vary in glyphosate resistance with and without drought stress.

|  | Population | Slope<sup>ab</sup> | Upper limit<sup>ab</sup> | GR<sub>50</sub><sup>ab</sup> | RI<sup>c</sup> |
| --- | --- | --- | --- | --- | --- |
| Dry Weight | RR WW | 0.61 ± 0.17 | 101.09 ± 6.40 | 22.31 ± 9.20 | 44.09 |
|  | RR DS | 0.05 ± 0.17 | 103.07 ± 6.13 | 47.39 ± 29.62 | 93.65 |
|  | R WW | 0.55 ± 0.09 | 100.99 ± 5.40 | 5.03 ± 1.40 | 9.94 |
|  | R DS | 0.48 ± 0.11 | 101.11 ± 5.30 | 19.70 ± 7.90 | 38.93 |
|  | S WW | 0.28 ± 0.12 | 81.60 ± 0.06 | 0.22 ± 0.30 | - |
|  | S DS | 8.60 X 10<sup>-5</sup> | 123.00 ± 0.01 | 0.50 ± 0.80 | 2.23 |

<sup>a</sup>Y = d/1 + exp[b(log x - log e)], Where: Y represents seedling dry matter at herbicide treatment rate x b, d, and GR50 represent the slope, upper limit, and herbicide rate that reduces seedling dry matter by 50%, respectively
<sup>b</sup> Mean ± SE
<sup>c</sup> RI refers to resistance index where the GR<sub>50</sub>/ GR<sub>50</sub>(SWW)

Glyphosate absorption of susceptible, resistant and highly resistant lines, both well-watered and after drought recovery, was modelled over time by asymptotic regression according to (Kniss *et al*., 2011), using the maximum absorption parameter (*A*_max_) and the time to reach 90% of maximum absorption (*t*_90_) to compare between populations and drought stress. Drought stress significantly reduced maximum absorption of glyphosate (*A*_max_) for all three populations, causing a minimum of a 45% (*P* < 0.01) reduction in glyphosate absorption; however, there were no statistically significant differences between population under well-watered or drought stress (Fig 3). The maximum glyphosate absorption in plants before drought stress ranged between 70% and 89% of applied glyphosate where maximum absorption after drought stress ranged between 36% and 46% of applied glyphosate (Fig 3). Glyphosate absorption rapidly increased from 1-24 HAT both before and after drought stress, yet after drought stress glyphosate absorption plateaus at 24 HAT where in well-watered condition glyphosate absorption plateaus at 72 HAT. There were no statistical differences in time to reach 90% of maximum absorption (*t*_90_) between populations and drought stress. Additionally, there was no significant difference found in glyphosate translocation after drought stress (Supplemental data 1). The relationship between periodic drought stress and reduced glyphosate absorption has previously been described in other weedy species such as *Amaranthus tuberculatus, C. truncata*, *S. oleraceus*, and *C. bonariensis* (Skelton *et al*., 2016; Peerzada *et al*., 2021). It’s well documented that under drought stress, plants respond by increasing the size and chemical composition of the cuticle, and that cuticle thickness and composition can have major impacts on glyphosate absorption (Trezzi *et al*., 2020).

**Figure 3.**
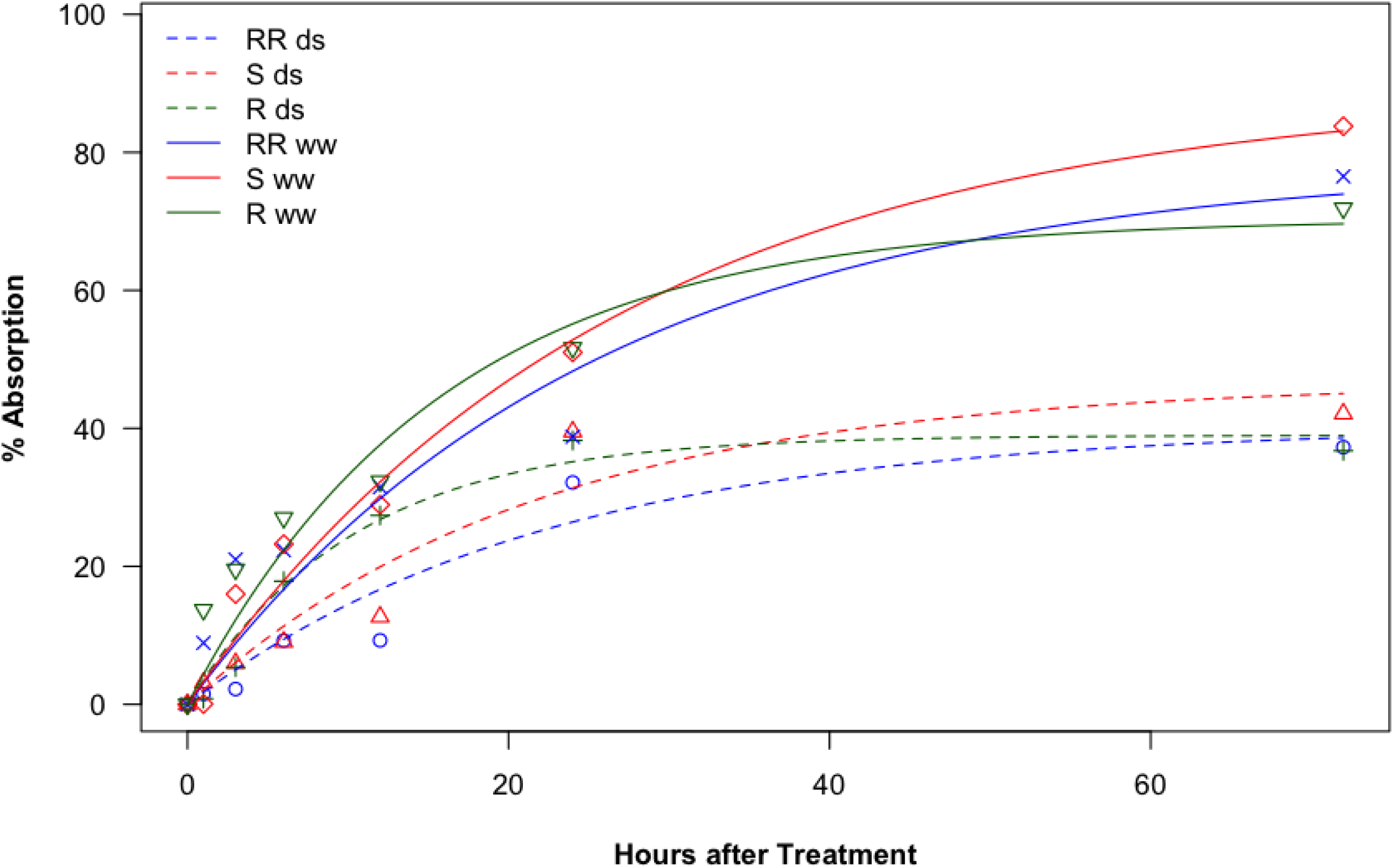
^14^C Glyphosate absorption assay *Erigeron canadensis* biotypes RR (highly resistant), R(resistant), and S(susceptible) before and after drought stress. Absorption is expressed as a percentage of applied. ds refers to drought stress, ds refers to well water.

### Gene co-expression network analysis reveals differences related to genotype, herbicide resistance, and drought stress

To understand how herbicide and drought stress impact horseweed at the cellular level, we performed a weighted gene co-expression network analysis (WGCNA) on RNAseq data collected from 1) well-watered, 2) glyphosate treated, 3) drought stressed, 4) drought stressed and recovered, and 5) drought stressed-recovered-glyphosate treated plants from S, R and RR populations. WGCNA identified 30 gene expression modules, where each module is represented by a module eigengene, which is an idealized gene which represents the overall gene expression trend in the module (Fig 4). The number of genes in each module ranged from 52 to 6091 (Fig 4a). Each module either had a positive or negative correlation to a treatment. While most modules show no strong correlation, we chose to take a detailed look at 3 modules that show expression differences that are correlated with populations, treatments, or populations x treatments.

**Figure 4.**
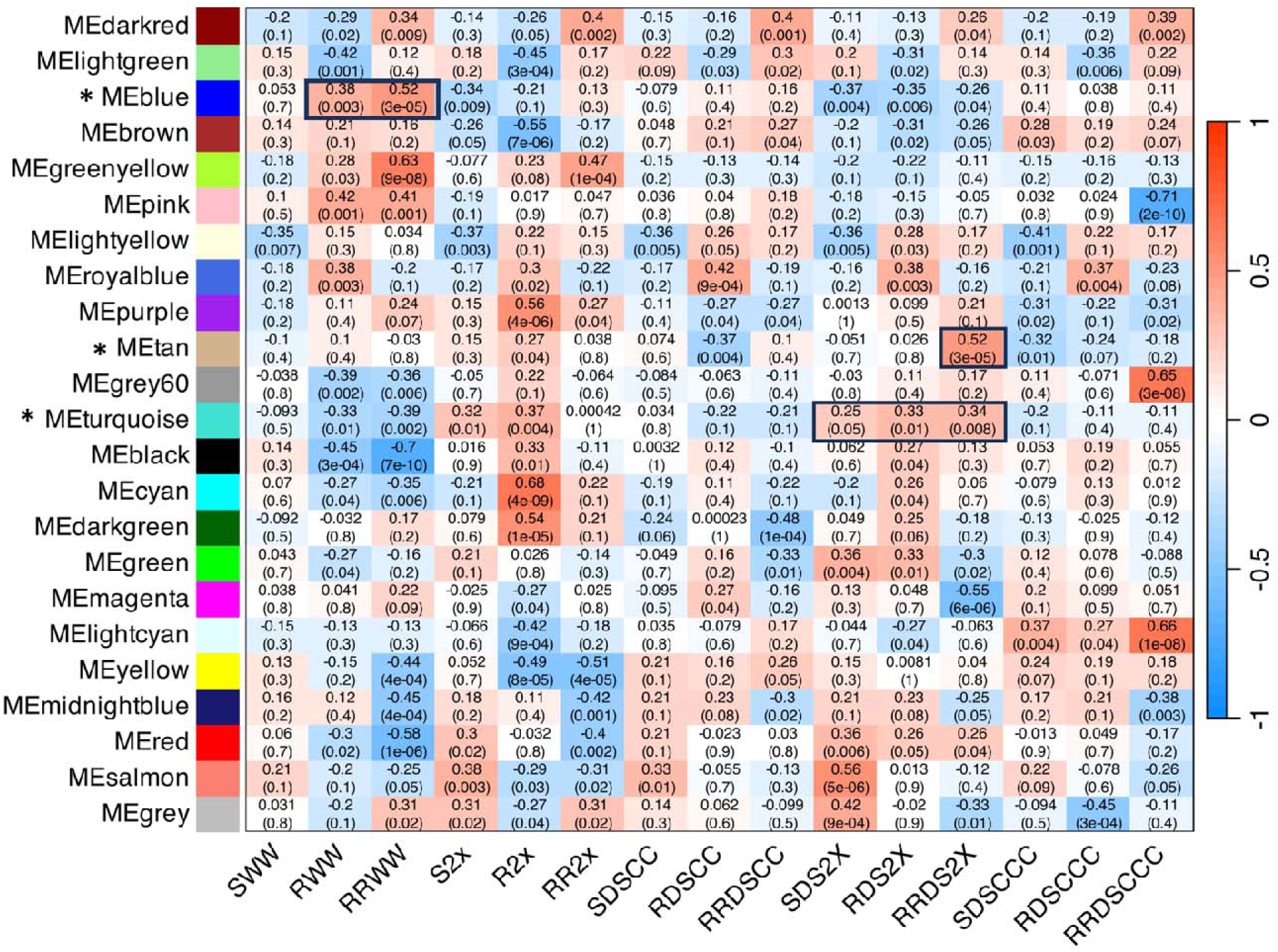
(A) Weighted gene co-expression heatmap. Identification of modules associated with either population or treatment. The upper and lower values represent Pearson correlation coefficient (PCC) and P value. The scale corresponds to the Pearson correlation coefficient (PCC). Modules with a (*) are discussed in the results. Significant correlations discussed in the results are highlighted with a box. RR (highly resistant), R(resistant), and S(susceptible), DS refers to drought stress, WW refers to well water. WW C refers to well watered controls, WW 2X refers to well plus glyphosate, DS CC refers to drought stress, DS CCC refers to drought plus recovery, DS 2X refers to drought stress plus glyphosate.

ME-Blue was correlated (p>0.001) with the R and RR populations under well-watered conditions (Fig 4). In this module we expect to find genes that are constitutively different in R compared to S populations (i.e. evolved resistance mechanisms). This module contained numerous genes related directly to glyphosate including overexpression of the glyphosate’s target protein EPSPS as well as other genes in the shikimate pathway such as shikimate O-hydroxycinnamoyl transferase and caffeoyl shikimate esterase, key enzymes for lignin biosynthesis. However, this module was large (5273 genes) and most likely contained numerous genes related more to population level differences that are not related to glyphosate as the populations were genetically distinct and horseweed has high levels of genetic diversity. A Gene Ontology (GO) term enrichment analysis of ME-Blue was conducted to identify the top GO terms, in which 30 GO terms were enriched including GO terms relating to carbohydrate metabolism, photosystem II assembly, and lipid biosynthesis (Fig 5a). Within lipid biosynthesis, we identified numerous genes related to the production of very long chain fatty acids (VLCFAs) and cuticle development.6/16/2026 1:23:00 PM This module contains seven 3-ketoacyl-CoA synthases genes that are involved in fatty acid elongation to produce Very Long Chain Fatty Acids, key components of cuticles; as well numerous genes central to cuticle development such as CER1 and MAH1 which lead to the production of epicuticle waxes (Joubès and Domergue, 2018). The gene CYP 86A8 was also included in this module which for producing hydroxylated fatty acids needed for cutin formation (Höfer *et al*., 2008). Additionally, this module contains multiple glycerol-3-phosphate acyltransferase (GPATs) responsible for the cellular transfer of cutin monomers and ABCG11 a known transporter of cutin and wax monomer out of the cell (Bird *et al*., 2007*a*). Therefore, there are most likely population level-differences in cuticle amount and composition that may factor into standing glyphosate resistance in these populations.

**Figure 5:**
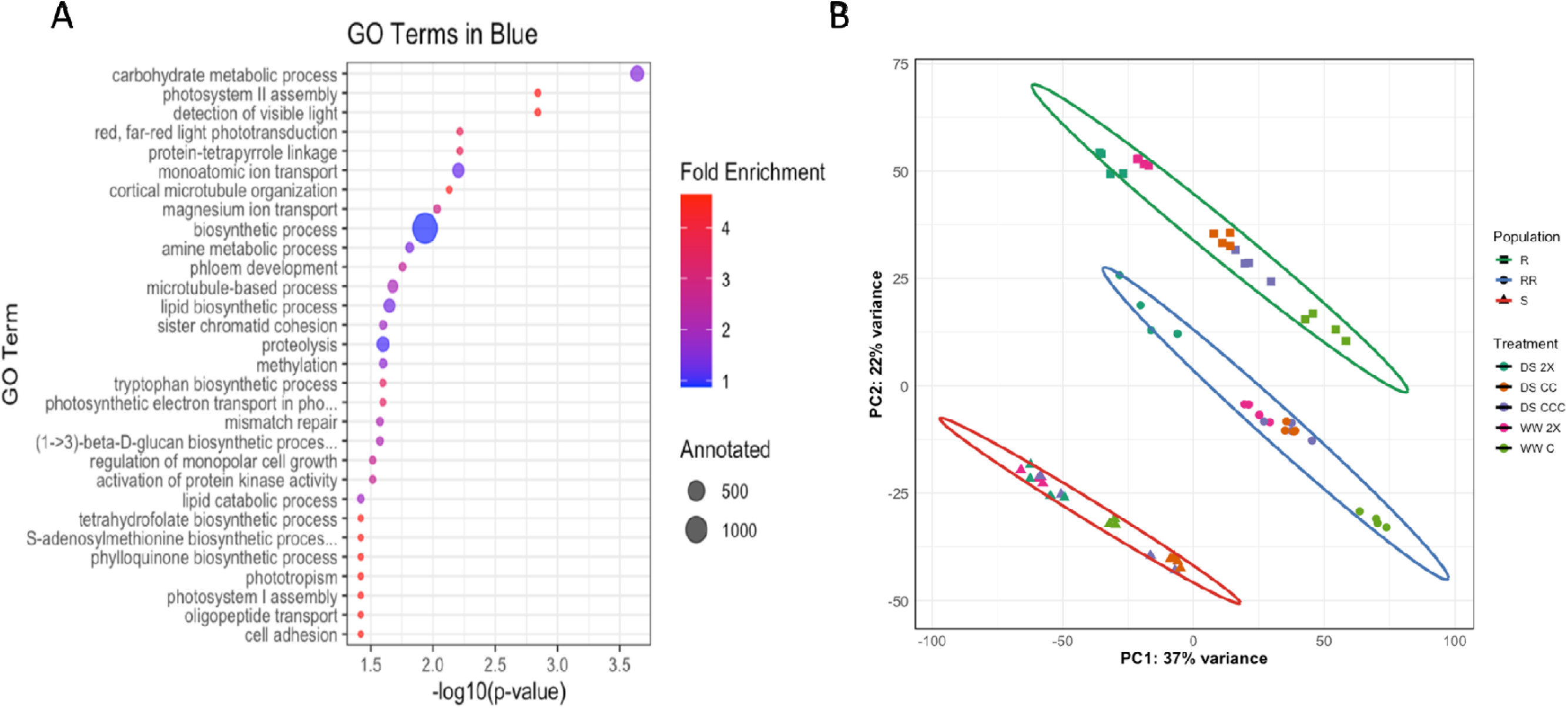
(A) GO term enrichment analysis of most enriched GO terms within Blue. -log10(FDR) is the -log10 of the false discovery rate adjusted false discovery rate p value. (B) Principal component analysis of gene expression during both drought and glyphosate stress. Ellipses represents 95% confidence intervals for each population. WW C refers to well watered controls, WW 2X refers to well plus glyphosate, DS CC refers to drought stress, DS CCC refers to drought plus recovery, DS 2X refers to drought stress plus glyphosate. RR (highly resistant), R(resistant), and S(susceptible), DS refers to drought stress, WW refers to well water.

The turquoise module correlated with combined drought and glyphosate application in all three populations, revealing how these combined stresses affect horseweed despite population differences. A Principal Component Analysis (PCA) revealed that under drought stress all populations cluster together and when glyphosate is applied there is a global shift in the PCA in all samples in the same direction (Fig 5b). This module captures gene expression trends associated with this shift according to the PCA plot. Within this module we found 5 different *Aldo-keto Reductases* (*AKR*) including an *Aldo-keto reductase* 3 and 5 which are overexpressed in both R and RR pops under glyphosate & glyphosate plus drought. AKRs have been shown to directly metabolize glyphosate leading to resistance in grasses and might partially explain increases in glyphosate tolerance after drought and glyphosate application, especially in the R and RR populations (Pan *et al*., 2019). Furthermore, multiple *ABCC transporters* are present in the turquoise module including *ABCC* 3,8,10,12,14, and 15. These genes can be critical genes in glyphosate metabolism; ABCC8 has been confirmed to be a vacuolar transporter of glyphosate in *Echinochloa colona* (Pan *et al*., 2021)*. ABCC* 10,12, and 14 all have been shown to be upregulated in glyphosate-resistant horseweed by others (Caygill and Dolan, 2023). Finally, multiple *UDP-glucuronosyltransferase* (UGTs) are present in the turquoise module such as 85C1, 85C2, 76B1 and 73E1 which have been found to be upregulated in glyphosate-resistant horseweed in the past (Caygill and Dolan, 2023). Recent work has shown that glycosylation of glyphosate mediated by UGTs can confer glyphosate resistance in rice (Yang *et al*., 2026). AKRs, UGTs and ABC transporters being co-expressed suggest that altered glyphosate metabolism, sequestration and translocation may all play a role in enhanced resistance after drought, and may partially explain the physiological results in this work.

Another interesting set of genes from the turquoise module relates to wax production and cuticle development, namely *WDS1*, *FAO* and certain *CYPs*. *WSD1* encodes a Bifunctional wax ester synthase/diacylglycerol acyltransferase, responsible for producing wax esters which is an ester of a fatty acid and a primary alcohol, helping to reduce cuticular transpiration (Li *et al*., 2008). *CYP94A2* encodes a cytochrome P450 capable of in-chain and terminal methyl hydroxylation involved in cutin formation (Pinot and Beisson, 2011). Additionally, this module contains multiple β*-amyrin 28-monooxygenases* which catalyze the carboxylation of β-amyrin at the C-28 position to form oleanolate (Han *et al*., 2013). It has been previously shown that when β-amyrin synthetase is overexpressed causing the overabundance of β-amyrin it led to increased cuticular water loss (Buschhaus and Jetter, 2012). These genes being co-expressed suggest that in response to glyphosate and drought stress, horseweed responds by increasing cuticle hydrophobicity potentially causing the reduction in glyphosate absorption after drought stress, as demonstrated by the absorption results in this manuscript.

Finally in the tan module we see genes that correlate with RR population under drought stress plus glyphosate stress. A GO term enrichment analysis revealed that this module is enriched for GO terms related to flowering and photoperiodism. One of the key genes identified was *GIGANTEA* (GI), a nuclear protein which is a major mediator between circadian clock genes and regulator of photoperiodic flowering time (Brandoli *et al*., 2020). GI regulates time to flowering via CONSTANS (CO) genes that control the expression of flowering time (FT). Two CONSTANS-like genes are also present in the tan module suggesting that they are being co-expressed with GI. In horseweed, it has been shown that when horseweed begins to bolt and flower, they become more resistant to glyphosate (Schramski *et al*., 2021; Fisher *et al*., 2023). The relationship between developmental stage and glyphosate resistance is still unclear, but these genes may play a key developmental role in both.

### Leaf Cuticular Waxes of Horseweed

To further investigate the involvement of the cuticle and the genes that regulate it, we analyzed the biochemical composition and quantity of epicuticular waxes of young leaves from the three horseweed populations, both before and after drought stress via GCMS. A total of 35 wax compounds were identified by comparing their mass spectra to those of authentic standards (Table 2). Among these, ten compounds were identified as alkanes, with chain lengths ranging from C25 (pentacosane) to C35 (pentatriacontane) (Table 2). In all three populations, hentriacontane (C31) was the predominant alkane (Table 2). Six compounds were identified as primary alcohols, with chain lengths ranging from C24 (tetracosanol) to C34 (dotriacontanol), with dotriacontanol being the most abundant primary alcohol in all three populations (Table 2). Nine compounds were identified as fatty acids, with chain lengths ranging from C21 (nonadecanoic acid) to C36 (Tetratriacontanoic acid) (Table 2). Two compounds were identified as aldehydes, specifically C30 and C32 aldehydes (Table 2). Additionally, six pentacyclic triterpenoids, namely β and α- amyrin and one sterol, were identified among the non-aliphatic compounds (Table 3).

**Table 2.**
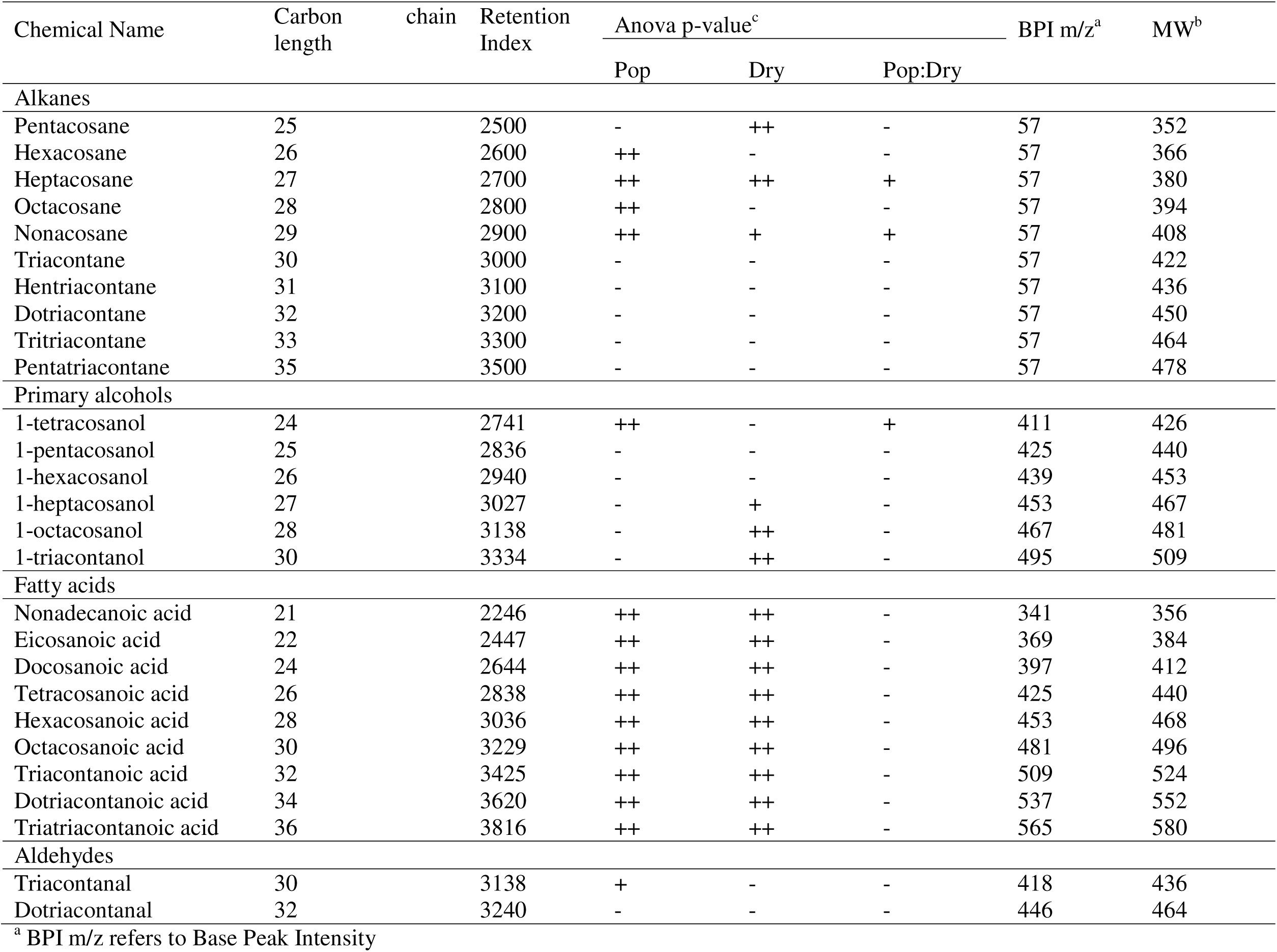
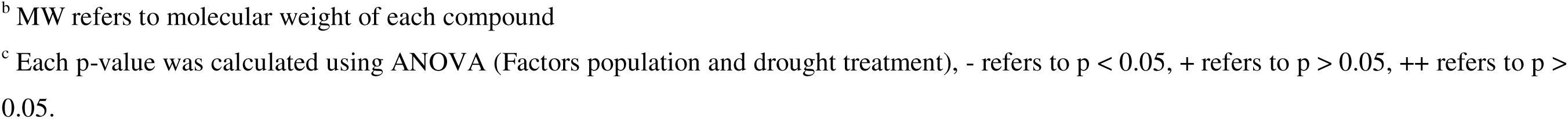
Chemical composition of aliphatic cuticular waxes detected on *Erigeron canadensis* leaves.

**Table 3.** Chemical composition of aliphatic cuticular waxes detected on *Erigeron canadensis* leaves.

| Chemical Name | Chemical class | Anova p-value<sup>c</sup> |  |  | BPI m/z<sup>a</sup> | MW<sup>b</sup> |
| --- | --- | --- | --- | --- | --- | --- |
|  |  | Pop | Dry | Pop:Dry |  |  |
| Erythrodiol | Triterpenoid | ++ | ++ | ++ | 218 | 442 |
| Sterol | Sterol | ++ | ++ | ++ | 329 |  |
| β-amyrin | Triterpenoid | ++ | ++ | ++ | 218 | 426 |
| β-amyrin acetate | Triterpenoid | ++ | ++ | ++ | 218 | 468 |
| α-amyrin | Triterpenoid | ++ | ++ | ++ | 218 | 426 |
| α-amyrin acetate | Triterpenoid | ++ | ++ | ++ | 218 | 468 |
| Oleanolic acid | Triterpenoid | ++ | ++ | ++ | 203 | 456 |

<sup>a</sup> BPI m/z refers to Base Peak Intensity
<sup>b</sup> MW refers to molecular weight of each compound
<sup>c</sup> Each p-value was calculated using ANOVA (Factors population and drought treatment), - refers to $p < 0.05$ , + refers to $p > 0.05$ , ++ refers to $p > 0.05$

Under well-watered conditions, the RR population contained the highest wax load with 24 ug/cm of wax, followed by R and S population with 20.6 and 20.3 ug/cm^2^ of wax respectively (Fig 6a). In terms of wax composition: alkanes 27-40%, primary alcohols 18-23% and cyclic triterpenoids 30-46% comprise most of the wax in all three populations (Fig 6b). In the RR population under well-watered conditions alkanes are the most abundant wax species comprising 40.6% of total wax, followed by triterpenoids 23.1% and primary alcohols 21.4% (Fig 6b). In both the R and S populations under well-watered conditions triterpenoids comprising 35.0% and 35.1% of total wax respectively, followed by alkanes (27.6% and 31.7%) and primary alcohols (23.6% and 18.2%) (Fig 6b). Among the minor components of waxes which were present on all three populations, were fatty acids (1.3–4.4%), aldehydes (0.8–2.7%) and sterols (7.6–11.3%) which make up most of the rest of the cuticle (Fig 6b).

**Figure 6.**
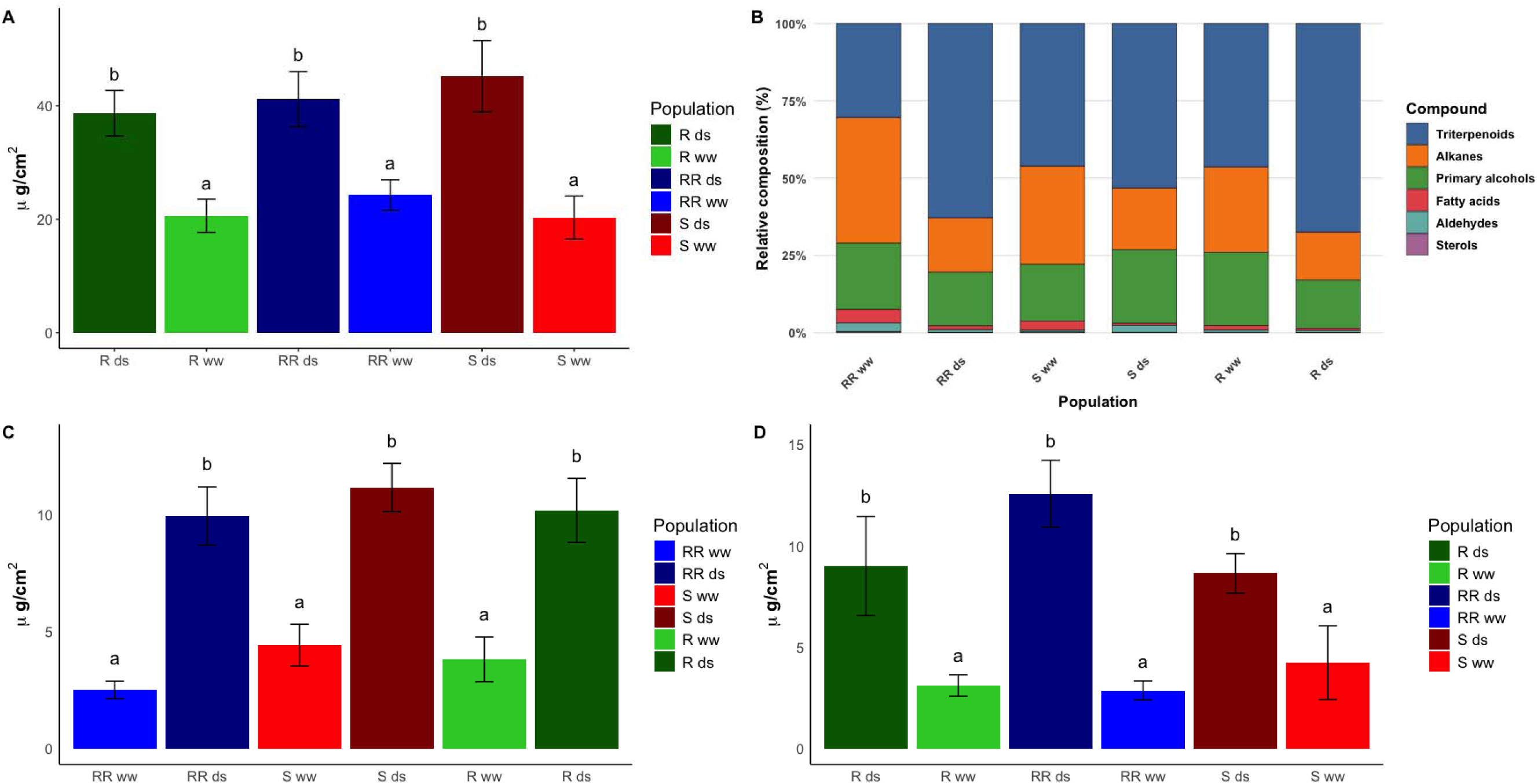
(A) Total wax content of cuticle waxes in *Erigeron canadensis* biotypes RR (highly resistant), R(resistant), and S(susceptible), before (ww) and after drought stress (ds). (B) Relative composition of cuticle waxes in *Erigeron canadensis* biotypes RR, R, and S before and after drought stress sorted by chemical class. (C & D) Concentrations of α and β amyrin in cuticle waxes in *Erigeron canadensis* biotypes RR, R, and S before and after drought stress.

When comparing total wax loads of plants grown under well watered and drought stressed conditions, we found that drought stress significantly increased wax loads for all three populations: in RR, drought stress increased wax load by 41.1%; in R, by 55.1%; and in S, by 46.7% (Fig 6a). For all three populations we found that the total amount of triterpenoids was mainly responsible for this increase in total wax load (Fig 6b). Under drought stress both RR and R population showed dramatic increases in triterpenoids (Fig 6b). RR went from 23.1% in well-watered conditions to 55.7% in drought stress, R went from 35.1% to 57.9%, and S went from 35.0% to 45.8% triterpenoids (Fig 6b). Of the four pentacyclic triterpenoids β and α- amyrin were most significantly induced by drought in all populations (RR 2.88 ug/cm^2^ to 12.58 ug/cm^2^, R 3.12 ug/cm^2^ to 11.55 ug/ cm^2^ and S 2.52 ug/cm^2^ to 9.16 ug/cm^2^ for β amyrin) and (RR 2.52 ug/cm^2^ vs 9.94 ug/cm^2^, R 3.822 ug/cm^2^ vs 10.11 ug/cm^2^ and S 4.44 ug/cm^2^ vs 11.1 ug/cm^2^ for α- amyrin). Additionally, erythrodiol, another triterpenoid, was induced under drought in both S (.005 ug/cm^2^ vs .014 ug/cm^2^) and R (.003 ug/cm^2^ vs .020 ug/cm^2^) but not in RR Under drought stress there was also an increase in the total amount of primary alcohols for all three populations. Unlike triterpenoids, the S population had dramatic increases in primary alcohols S 3.71 ug/cm^2^ vs 10.76 ug/cm^2^ compared with RR 5.19 ug/cm^2^ vs 7.13 ug/cm^2^ and R ug/cm^2^ vs 11.55 ug/cm^2^. Dotriacontanol was the dominant primary alcohol for all three populations and was significantly induced after drought stress in both RR and S populations. In the RR population in particular there was a significant decrease in VLCFAs, namely triatriacontanoic acid, dotriacontanoic acid, and triacontanoic acid. In summary, all three populations had similar changes to their wax profile after drought stress, dominated by increases in β and α- amyrin.

### Genes responsible for cuticle composition following drought

Following drought stress, there were substantial changes both in the cuticle composition and the gene expression of cuticle-related genes. There was a reduction in Very Long Chain Fatty Acids (VLCFAs) following drought stress (Fig 6b), dotriacontanoic acid, and triacontanoic acid in the RR population (Fig 7a). Simultaneously, there were increases in primary alcohols, especially dotriacontanol and triacontanol. During cuticle biosynthesis, VLCFAs are converted into primary alcohols by fatty acyl-CoA reductases (i.e. *CER4* genes). CER4 was overexpressed following drought stress in RR populations (Fig 7b). This overexpression is the most likely cause of the observed VLCFA-Primary alcohol shift in cuticle composition in response to drought (Rowland *et al*., 2006). *3-oxoacyl synthase* (KAS) was overexpressed following drought stress across all populations. This gene catalyzes the condensation of acyl-ACP with a malonyl-ACP to elongate the fatty acid carbon chains which can be further elongated into VLCFAs by KCS enzymes (Zhou *et al*., 2020). Overexpression of KAS creates the building blocks for all cuticle aliphatic compounds. Cuticle lipids are exported out of the cell into the extracellular matrix ABC transporters such as ABCG11 (Bird *et al*., 2007*b*), ABCG11 was drought induced in both resistance populations (Fig 7b). Taken together, plants following drought stress modulate gene expression leading to changes in aliphatic compounds.

**Figure 7.**
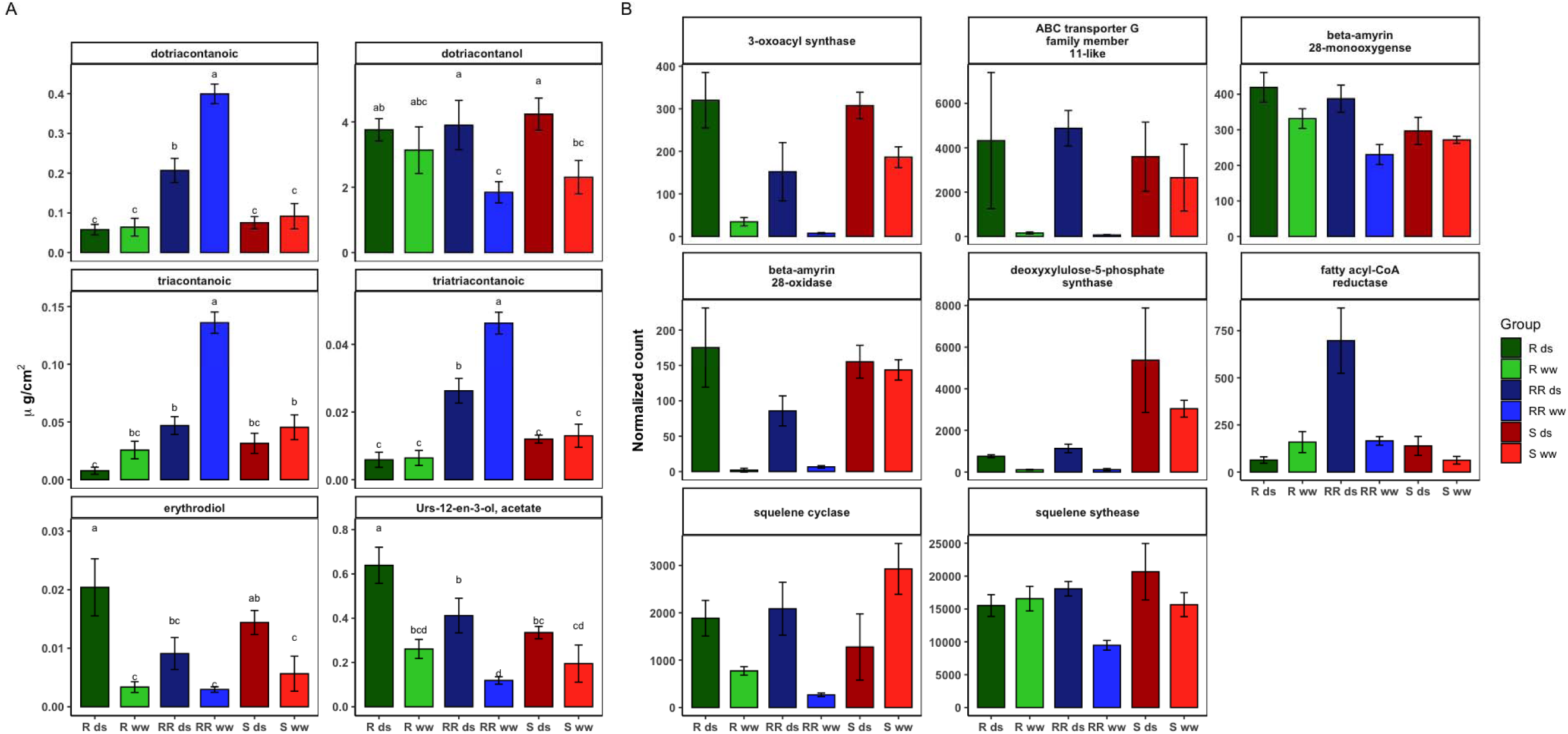
Cuticle metabolites and gene expression across populations and drought treatments. Bar plots (mean ± SE) show relative metabolite abundance (A) and normalized gene expression (B) across population × treatment groups (RR_w, RR_ds, S_w, S_ds, R_w, R_ds). Letters denote significant differences among groups (P < 0.05).

The largest changes in cuticle composition after drought was in pentacyclic triterpenoids. Several triterpenoid biosynthesis genes were upregulated following drought, specifically *Deoxyxylulose-5-phosphsate synthase*, β*-amyrin 28-monooxygenases* and *oxidase*, and *Squalene synthase* and *Squalene cyclase*. Deoxyxylulose-5-phosphsate synthase (DXS) is the rate-limiting enzyme controlling the entry of carbon into the MEP pathway, one to two pathways producing isoprene subunits key precursors of triterpenoid (Gawriljuk *et al*., 2025). Squalene synthase (SQS) catalyzes the condensation of two farnesyl pyrophosphate molecules into squalene (Liu and Fu, 2018). Squalene cyclases are the rate-limiting enzymes in triterpenoid biosynthesis, catalyzing the conversion of squalene into 2,3-oxidosqualene (Kumar *et al*., 2025), the precursor molecule for α and β amyrin. Overexpression of several rate-limiting genes in both terpenoid and triterpenoid biosynthesis most likely the primary driver of the nearly 300% increase in both α- and β-amyrin. These genes were also overexpressed when compared to β-amyrin synthase, over production β amyrin is being driven earlier in the synthesis pathway rather than at the terminal reaction step, β-amyrin synthesis. In addition to straight increase in α- and β-amyrin levels, multiple triterpenoids such as α-amyrin acetate (Urs-12-en-3-ol acetate), erythrodiol and oleanolic acid levels also increased following drought. The conversion from β amyrin to oleanane is facilitated by β-amyrin 28-monooxygenases which specifically catalyze the carboxylation of β-amyrin at the C-28 position to first form erythrodiol and then oleanolic acid.

In summary, cuticle composition changes in all three populations in response to drought can be explained by: 1) overexpression of CER4 shifts the composition from VLCFAs to more primary alcohols.2) Overexpression of *Deoxyxylulose-5-phosphsate synthase*, β*-amyrin 28-monooxygenases* and *oxidase*, and *Squalene synthase* and *Squalene cyclase* leads to the massively increased α and β amyrin content, and3) increased β-amyrin 28-monooxygenases leads to increased erythrodiol and then oleanolic acid concentration. These changes are primarily occurring to thicken the cuticle and make it less permeable to water, to prevent additional water loss during drought stress. However, this decreased permeability goes both ways and also decreases water absorption across the leaf surface, an essential process in foliar applied herbicide application.

### The role of the cuticle in glyphosate absorption

The polar path of diffusion refers to the way polar molecules, including organic ions like glyphosate, diffuse through the cuticle. Unlike other polar molecules, water can diffuse across the plant cuticle via both lipophilic cutin and wax, owing to its high mobility and small molecular size (Schreiber, 2005). Several herbicides, such as glyphosate, are weak acids and are charged (Shaner, 2010). These ions are not soluble in the lipophilic cutin and wax fractions of the cuticle because they have a hydration shell (Stein, 1967). Yet glyphosate still diffuses across the cuticle; therefore, it is hypothesized that polar, charged molecules can diffuse across the cuticle via aqueous pores or via mechanical damage in the cuticle (Schreiber, 2005). The exact nature of cuticle aqueous pores remains unknown, but these pores are transient structures that depend on the cuticle hydration level (Schreiber, 2005). Cuticle wax contains polar functional groups, such as primary alcohols and fatty acids, that can absorb water. It’s possible that these aqueous pores form as polar functional groups create hydration channels in the cuticle, allowing charged herbicides to diffuse across the cuticle. Many research groups have tried to locate aqueous pores in the cuticle using AgNO_3_ staining. This works because Ag ions precipitate out of solution in the apoplast, creating a dark stain (Schreiber, 2005). AgNO_3_ staining shows that aqueous pores are primarily located around the base of trichomes, at cell edges, and around stomata (Schreiber, 2005). Following drought stress, there was a substantial increase in all four pentacyclic triterpenoids present in the cuticle, including α- and β amyrin, ursane, and oleanane. These pentacyclic triterpenoids accumulate within the intracuticle region and serve as an interface between epicuticle wax and cutin. The role of cuticular triterpenoids is mainly to improve mechanical strength, reinforce the cutin monomers, and serve as structural fillers within the cutin matrix. Due to glyphosate’s highly polar nature, it is unable to diffuse through non-polar waxes, but it can diffuse through aqueous pores or through mechanical damage. Increases in cuticular triterpenoids after drought would lead to greater mechanical strength and filtering within the intracuticle region, possibly reducing glyphosate absorption following drought stress.

## Conclusions

Drought stress substantially reduced glyphosate sensitivity in horseweed across susceptible, resistant, and highly resistant populations, with the largest changes observed in both the resistant and highly resistant populations, indicating that drought acts as a significant resistance[enhancing environmental modifier. Phenotypically, drought stress significantly reduced maximum glyphosate absorption by at least 45% in all populations and accelerated the plateau of absorption, consistent with increased physical barriers to foliar uptake rather than altered translocation. Transcriptomic analyses supported coordinated changes in gene expression associated with both resistance and drought, including population-specific overexpression of genes involved in the shikimate pathway, lipid biosynthesis, and cuticle development (e.g., KCS, CER1, MAH1, CYP86A8, GPATs, and ABCG11), suggesting genetically encoded differences in cuticle formation among populations under well-watered conditions and become further differentiated after drought stress. Additionally, drought- and glyphosate-responsive modules showed co-expression of detoxification and transport genes (AKRs, UGTs, and ABCC transporters), indicating potential contributions of enhanced metabolism and sequestration to resistance following drought priming. Biochemical analyses of the cuticle confirmed that drought stress increased total epicuticular wax loads and altered wax composition in all populations, with pronounced increases in hydrophobic triterpenoids (notably β- and α-amyrin) and primary alcohols, changes that are consistent with reduced cuticular permeability.

Together, these results demonstrate that drought-induced phenotypic reductions in glyphosate absorption are mechanistically linked to both transcriptional upregulation of cuticle biosynthesis pathways and quantitative and qualitative changes in cuticular wax composition, providing a mechanistic framework for drought-mediated enhancement of glyphosate resistance in horseweed. This is most likely especially pronounced in a highly polar herbicide such as glyphosate which relies heavily on cracks and pores in the adaxial leaf surface for absorption, as it cannot penetrate even moderately hydrophobic barriers. Ultimately, this contributes to our understanding about the importance of environmental conditions, both at the time of application and in the weed’s history, for herbicide efficacy and supports the cuticle as a major integrator of stress resistance in weeds. Given what we know about the cuticle’s importance in water retention, it is highly likely to be involved in other stresses as well including drought and cold tolerance.

## Supplementary data

S1 Anova results for glyphosate translocation

## Data Availability

The RNA-seq data is available at PRJNA1433610.

## Acknowledgements

We would like to thank Dr. Tony Schilmiler and James O’Keefe from the MSU Mass Spectrometry Core. We also thank Jan Michaels for her assistance during this experiment. The second author, Ahmet Tansel Serim, also gratefully acknowledges the financial support given by the Scientific and Technical Research Council of Turkey (TUBITAK) for the 2219 International Postdoctoral Research Fellowship Programme (Grant No. 1059B192200396). We would also like to thank the Unites Soybean Board for providing funding for partial graduate student assistantship (Project No. 26-206-S-C-2-G).

## Author Contribution

MO,AT,EP: conceptualization; MO,AT,MM,EP: methodology; MO,AT,MM,EP: formal analysis; MO,AT,MM,EP: investigation; EP: resources; MO,AT,MM,EP: data curation; MO,AT,MM,EP: writing - original draft; MO,AT,MM,SAR,EP: writing - review & editing; MO,AT,MM,SAR,EP: visualization; and EP: funding acquisition

